# Template Switching Mediates Contractions of a Repetitive Coding Region in *S. cerevisiae*

**DOI:** 10.1101/712588

**Authors:** Taylor Stewart, Alexandra E. Exner, Paras Patnaik, Stephen M. Fuchs

## Abstract

The C-terminal domain (CTD) is an essential domain of the largest subunit of RNA polymerase II, Rpb1p, and is composed of 26 tandem repeats of a seven-amino acid sequence, YSPTSPS. Despite being an essential domain within an essential gene, we have previously demonstrated that the CTD coding region is genetically unstable. Furthermore, yeast with a truncated or mutated CTD sequence are capable of promoting spontaneous genetic expansion or contraction of this coding region to improve fitness. We investigated the mechanism by which the CTD contracts using a tet-off reporter system for *RPB1* to monitor genetic instability within the CTD coding region. We report that contractions require the post-replication repair factor Rad5p but, unlike expansions, not the homologous recombination factors Rad51p and Rad52p. Sequence analysis of contraction events reveals that deleted regions are flanked by microhomologies. We also find that G-quadruplex forming sequences predicted by the QGRS Mapper are enriched on the noncoding strand of the CTD compared to the body of *RPB1*. Formation of G-quadruplexes in the CTD coding region could block the replication fork, necessitating post-replication repair by template switching. We propose that contractions of the CTD result when microhomologies misalign during Rad5p-dependent template switching via fork reversal.

## INTRODUCTION

RNA polymerase II is an essential eukaryotic protein complex that is responsible for the transcription of mRNA. The largest subunit of this complex, Rpb1p, has a C-terminal domain (CTD) that serves as an essential binding domain for numerous transcription factors as well as proteins involved in chromatin remodeling and DNA repair (1, 2). The CTD is composed of tandem repeats of a seven-amino acid sequence, YSPTSPS. The number of repeats varies across organisms and generally increases with increasing organismal complexity: budding yeast typically have 26 repeats, while mammals can have up to 52. Studies have shown, however, that as few as eight repeats can support growth in yeast (Nonet et al. 1987). Despite being an essential region of an essential gene, we have demonstrated that the CTD coding region is genetically unstable. In addition to repeat length variation across organisms, previous work by our lab has shown that the CTD varies in length somewhat across strains of yeast, and yeast with a suboptimal CTD length or mutated CTD sequence are capable of promoting spontaneous expansion or contraction of the coding region in order to improve fitness (4). Repeat instability may therefore serve as a mechanism for reducing mutagenesis in an essential sequence by promoting the removal or templated repair of damaged repeats while maintaining overall length.

Tandem repeats are well known to be highly unstable, with mutation rates 10 to 100,000 times higher than the genomic average. Variable tandem repeats are found in a variety of genomic locations, including promoter regions as well as within coding regions, and tandem repeat instability is well known to be associated with disease (5–7). For example, trinucleotide repeats are known to be capable of undergoing expansions that can lead to a variety of neurodegenerative diseases. These repeats form stable hairpins that can lead to errors during DNA repair and replication (8). In addition to disease, there are numerous documented cases in which changes in repeat copy number in coding regions results in variable phenotypes that enable organisms to adapt to different environmental conditions (9–12). Tandem repeat instability may therefore be an important driver of evolution (7).

Unlike trinucleotide repeats, little is known about the mechanisms by which complex tandem repeats expand and contract. We are using the CTD, which is comprised of a degenerate 21-base pair repeat, as a model for complex tandem repeats. The CTD, as well as other complex tandem repeats, differs from the simpler trinucleotide repeats because it is not perfectly repetitive and therefore likely does not form canonical hairpin structures. Our lab has developed a tet-off reporter system to monitor expansions and contractions in the CTD, and genetic studies performed by our lab have revealed that homologous recombination factors are required for expansions but not contractions of the CTD. We therefore sought to determine the mechanism(s) by which the CTD undergoes contractions.

In this manuscript, we measured the frequency of contractions of the CTD in the absence of key DNA repair proteins and determined that contractions require the post-replication repair factor Rad5p but not Rad52p or Rad51p. We also analyzed the sequence of contraction events and found that microhomologies flank the repair junctions. Based on these findings, we propose that template switching via fork reversal mediates contractions of the variable CTD of RNA Polymerase II independently of homologous recombination factors.

## MATERIALS AND METHODS

### Yeast Strains and Plasmids

Strains used in this study were derived from GRY3019 (*MATa his3*Δ *leu2*Δ *lys2*Δ *met15*Δ *trp1*Δ::*hisG URA::CMV-tTA kanRPtetO7-TATA-RPB1*) (13). DNA repair mutants were constructed by heterologous gene replacement and verified by PCR with the primers in Supplemental Table 1. Yeast were grown on synthetic complete (SC) dropout medium or YPD at 30°C. Doxycycline (+DOX, 50 µg/ml) was added to plates to control the expression of genomic *RPB1* when appropriate. Plasmids used in this work were described previously in Morrill et al. 2016. Plasmids were freshly transformed into yeast and maintained on SC media lacking leucine (SC-Leu).

### Spotting Assays

For phenotypic growth assays, yeast expressing the appropriate plasmid were grown overnight in SC-Leu. Saturated overnight cultures were used to start fresh cultures in the same medium at an A_600_ of 0.2. The cells were allowed to double at least two times before approximately 1.0 × 107 cells were harvested and resuspended in sterile water in a 96-well plate. Cells were serially diluted ten-fold five times and then spotted onto SC-Leu plates with and without doxycycline using a 48-pin replicating tool. Plates were incubated at 30°C and imaged after three days.

### Suppressor Analysis

Individual colonies expressing p4stop were grown overnight in SC-Leu in a 96-well plate. Cultures were serially diluted ten-fold four times in sterile water, and each of the four dilutions were spotted onto SC-Leu+DOX plates. The range of dilutions ensured that single colonies for each culture could be identified for analysis by colony PCR. The plates were allowed to grow for three to four days, or until colonies were sufficiently large for colony PCR. Primers flanking the CTD were used to amplify the CTD coding region (Table S1). To ensure each suppressor is the result of an independent mutagenic event, only one colony per culture was analyzed. Colony PCR products were visualized by gel electrophoresis using 1% agarose in TBE and a subset of products were sequenced by traditional Sanger sequencing. Suppressors from at least three independent plasmid transformations were analyzed, and contraction frequencies were calculated from an aggregate of the total number of contractions observed relative to the total number of suppressors analyzed.

## RESULTS

### Repeat-length instability within the CTD of RNA Polymerase II

Others have studied the evolution of the repetitive CTD of RNA Polymerase II across species (14–17). Using budding yeast as a model, we set out to more closely examine how this sequence has evolved. Sequence alignments of the CTD from 93 *S. cerevisiae* genomes (18) revealed that while the total length of the CTD is generally conserved between 24 and 26 repeats, there have been multiple, independent rearrangements within the CTD repeat region (Figure 1 and Figure S1). This suggests that despite a strict requirement for conservation of its length, there is a history of both expansions and contractions within the CTD. Microhomologies throughout the sequence may serve as templates for these rearrangements.

**Figure 1.**
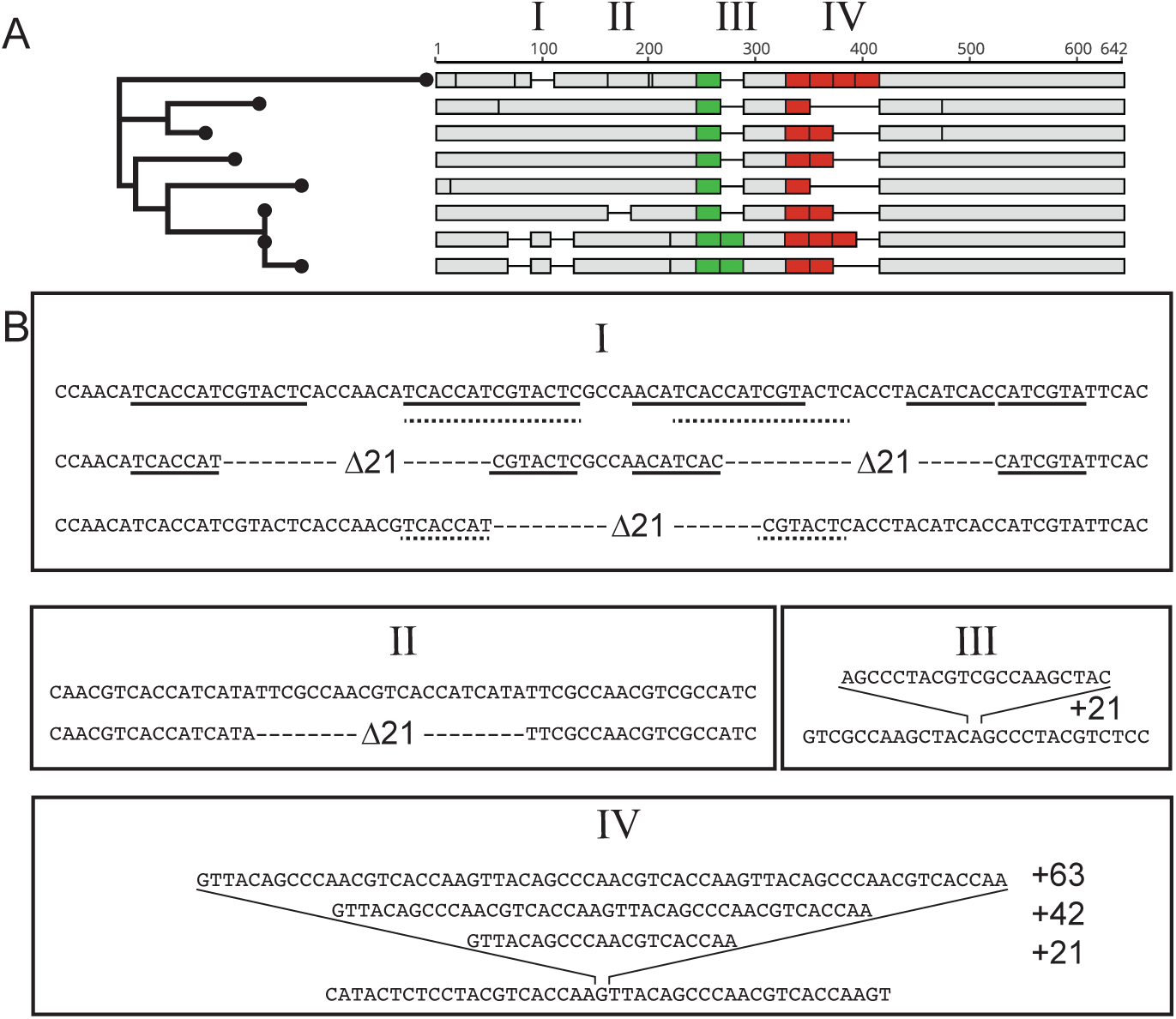
Sequence comparison of *S. cerevisiae* strains reveals multiple, independent instances of expansion within the repetitive C-terminal domain coding region. A) Sequence alignment of a subset of strains from the analysis by Strope et al. demonstrating four regions of expansion and contraction (I–IV). B) Analysis of the four regions showing that variable-length regions are flanked by regions of homology (underlined) and that all indels are some multiple of 21bp.

The repetitive nature of the CTD coding sequence likely enables it to rapidly evolve. While the mechanisms of repeat instability in simple repeat sequences, such as trinucleotide repeats, are well studied (8, 19), little is known about how more complex repeat sequences like the CTD promote variability. Our lab previously determined that expansions of the CTD require Rad52p, suggesting that expansions result from homologous recombination (4). In this work, we sought to determine the mechanism(s) by which the CTD contracts.

### CTD contraction frequency in the absence of key DNA repair proteins

We previously reported the mutation rate for cells expressing a plasmid-based mutant CTD construct designed to monitor contractions, p4stop (4). This construct encodes all 26 yeast CTD repeats, with repeats 8–11 each containing a stop codon (Tyr_1_→stop) and a non-coding Ser_2_→Trp mutation (Figure 2A). Previous studies have determined that a minimum of eight CTD repeats are required for efficient growth (20), and we have demonstrated that the protein produced from our 4stop mutant contains only seven CTD repeats (4).The expression of this construct is controlled by a tet-off system that we developed based on the work of Strathern and others (13). Briefly, the 4stop variant is under the control of the native *RPB1* promoter on a CEN/ARS-containing plasmid that is transformed into yeast. The genomic copy of *RPB1* is under the control of a tetracycline-responsive promoter, and doxycycline is added to the growth medium when appropriate in order to repress transcription of the genomic copy of *RPB1* (Figure 2B). Under these conditions, the cell must rely on the 4stop variant of Rpb1p for transcription. Thus, only cells that have acquired a spontaneous mutation that bypasses this selection, often through rearrangement of the *RPB1* coding sequence itself, grow on plates containing doxycycline (Figure 2C).

**Figure 2.**
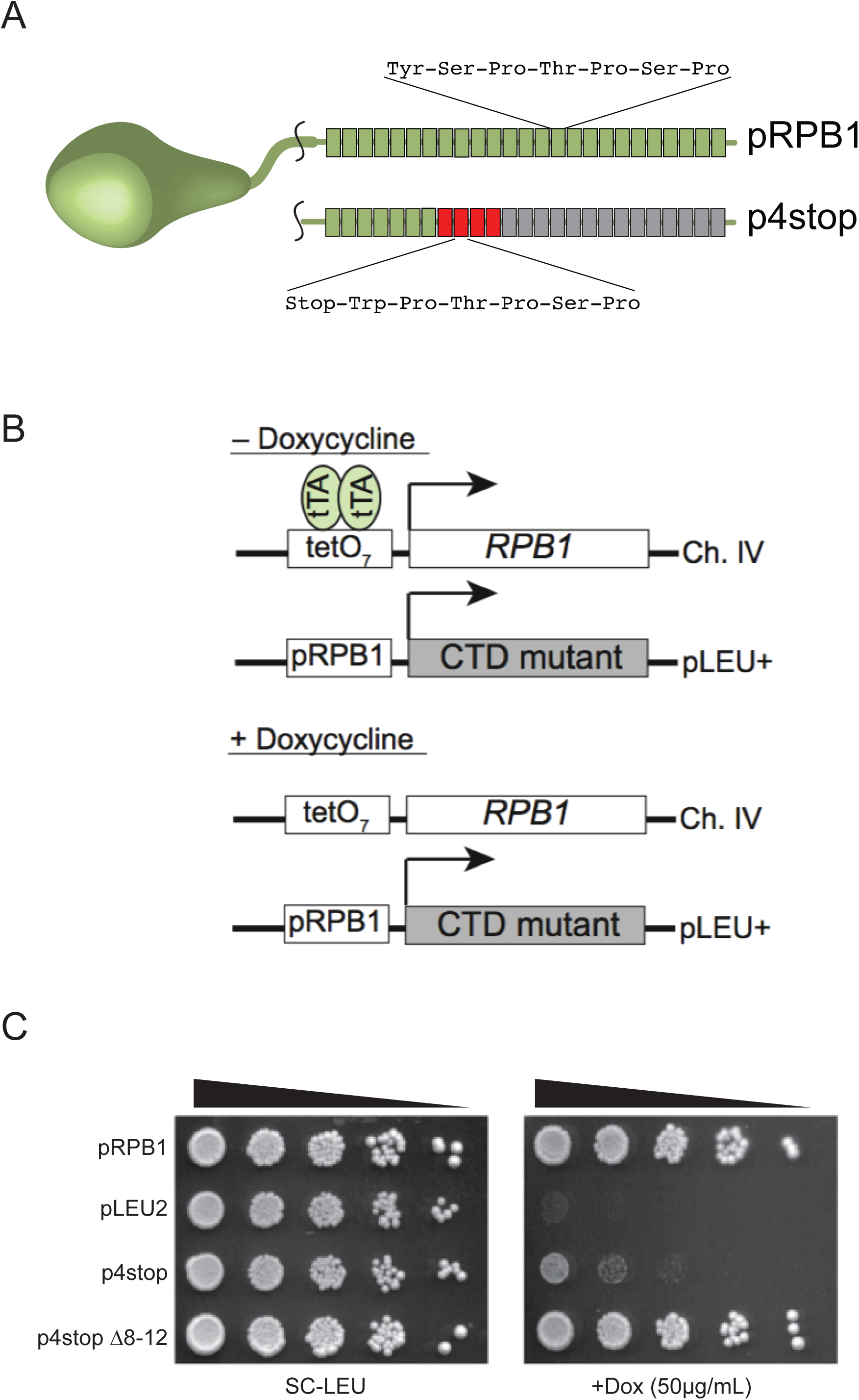
A genetic system for measuring changes in CTD repeat length. A) Schematic of the 4stop mutant CTD constructs. Each block represents one 21-bp repeat encoding the seven-amino acid CTD consensus sequence. pRPB1 encodes all 26 wild-type yeast CTD repeats, whereas p4stop encodes 26 repeats but is interrupted by four stop codons, indicated by red blocks. The protein produced from p4stop has only seven functional CTD repeats. B) Tet-Off system for monitoring contractions within the CTD. In the absence of doxycycline, the Tet transactivator (*tTA*) binds to tetO_7_ sites upstream of the genomic copy of *RPB1*, allowing transcription. In the presence of doxycycline, transcription of the genomic copy is repressed, and cells rely on a plasmid-based copy of *RPB1* under the control of its endogenous promoter. C) Spotting assay demonstrating the effectiveness of our Tet-Off system. pLEU2 is pRS315. p4stop produces a protein product that results in poor yeast viability in the presence of doxycycline. This phenotype is rescued by a contraction event that removes the four repeats containing stop codons. Spotting assays are representative examples of at least three independent trials.

We sought to determine the mechanism(s) through which the CTD contracts first by measuring the frequency of contractions in the absence of key DNA repair proteins. The p4stop plasmid was transformed into a series of isogenic strains in which different DNA repair genes were deleted, and cells were grown to large colonies to allow for the accumulation of spontaneous mutations. Cultures started from these colonies were then spotted on doxycycline to select for suppressors (Figure 3A). These fast-growing suppressors represent three types of events: 1) contractions within the plasmid that removed the four repeats containing stop codons, 2) homologous recombination events where the mutant plasmid copy of *RPB1* underwent a rearrangement with the doxycycline-regulated copy of *RPB1* in the genome, and 3) mutations elsewhere in the genome (4). The overall rate of suppressor formation, measured by fluctuation analysis, is similar in all mutants (Supplemental Table 2). In the present study, unique suppressors were characterized by colony PCR in order to determine the fraction of suppressors that were contractions.

**Figure 3.**
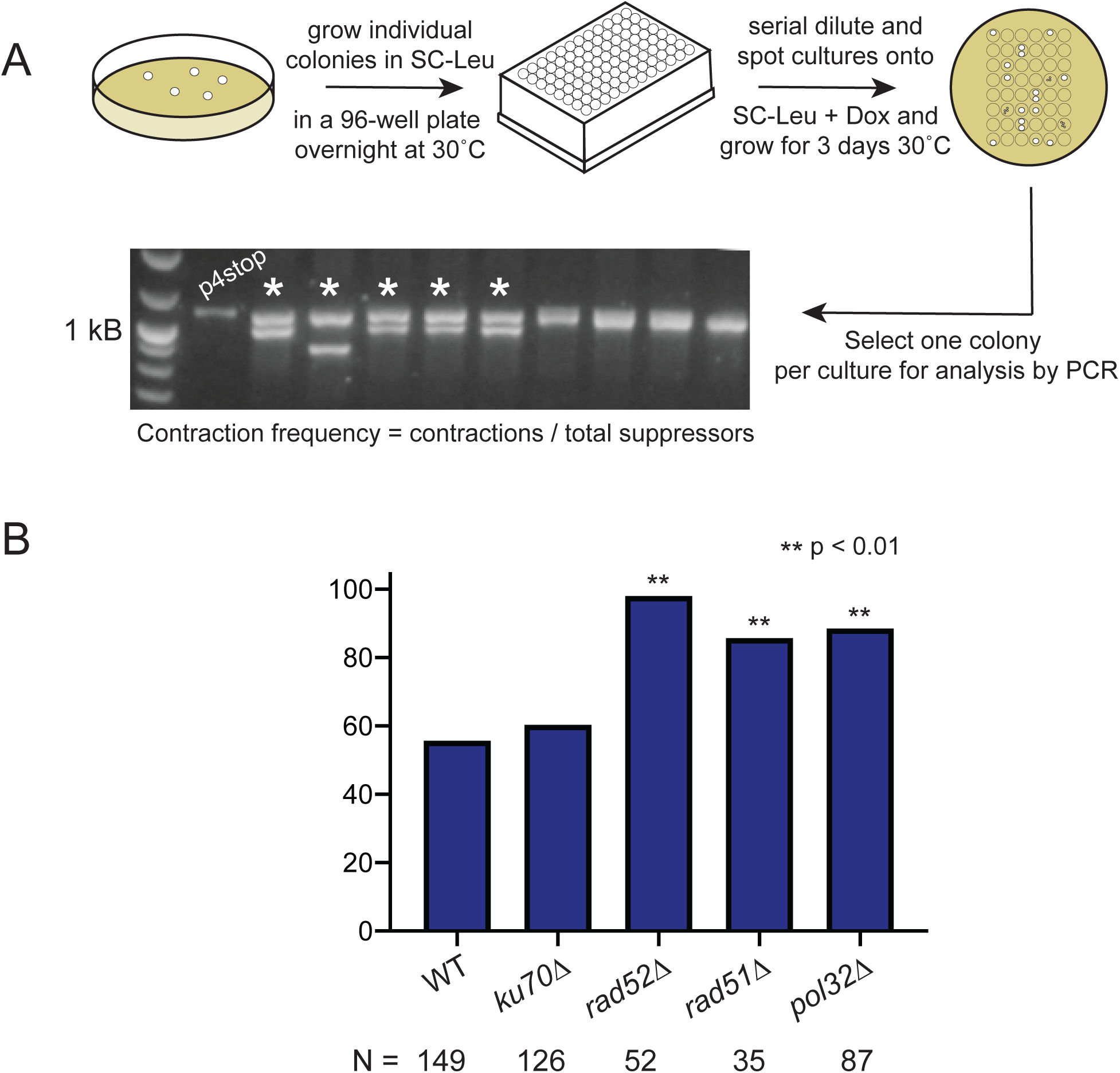
Analysis of CTD contraction frequencies. A) Suppressor generation and characterization assay. p4stop is transformed into yeast and maintained on SC-Leu, and individual colonies are used to start overnight cultures. Saturated overnight cultures are serially diluted and spotted on SC-Leu+DOX. One colony, representing a unique suppressor event, is selected from each spot for analysis by colony PCR. The contraction frequency is calculated by dividing the number of contractions observed by the total number of suppressors analyzed. B) Contraction frequencies of the CTD in the absence of key DNA repair proteins. Frequencies are determined by the accumulation of data from three independent plasmid transformations, and statistical significance was determined using a two-proportions z-test.

In the parent strain we find that 56 percent of fast-growing cells suppress the slow growth phenotype of 4stop by contraction within this region, removing the stop codons (Figure 3B). The frequency of contractions in a *ku70*Δ background is 60 percent, which is not significantly different from wild type based on a two-proportion Z test. However, in both *rad52*Δ and *rad51*Δ backgrounds, contractions occur significantly more frequently compared to wild type (98 and 86 percent, respectively). We similarly observed a high frequency of contractions (89 percent) in the *pol32*Δ background.

In addition to the primary DNA repair pathways, we also investigated the role of post-replication repair (PRR) in contractions. PRR consists of two pathways, Translesion Synthesis (TLS) and Template Switching (TS), both of which require Rad5p. In a *rad5*Δ mutant background, contractions of the CTD are almost completely abolished, defining PRR as a key pathway in spontaneous rearrangements within the CTD coding region (Figure 4A). In TLS, Rad5p physically interacts with Rev1p, which acts as a scaffold for translesion polymerases that bypass DNA lesions by inserting a base, potentially incorrectly, across from a lesion (21). While this pathway is often mutagenic, it generates point mutations and is unlikely to be a source of repeat-length instability. Nonetheless, to confirm that TLS does not play a role in contractions of the CTD, we measured the frequency of contractions in a *rev1*Δ background and found that the contraction frequency is not significantly different from wild type (Figure 4A).

**Figure 4.**
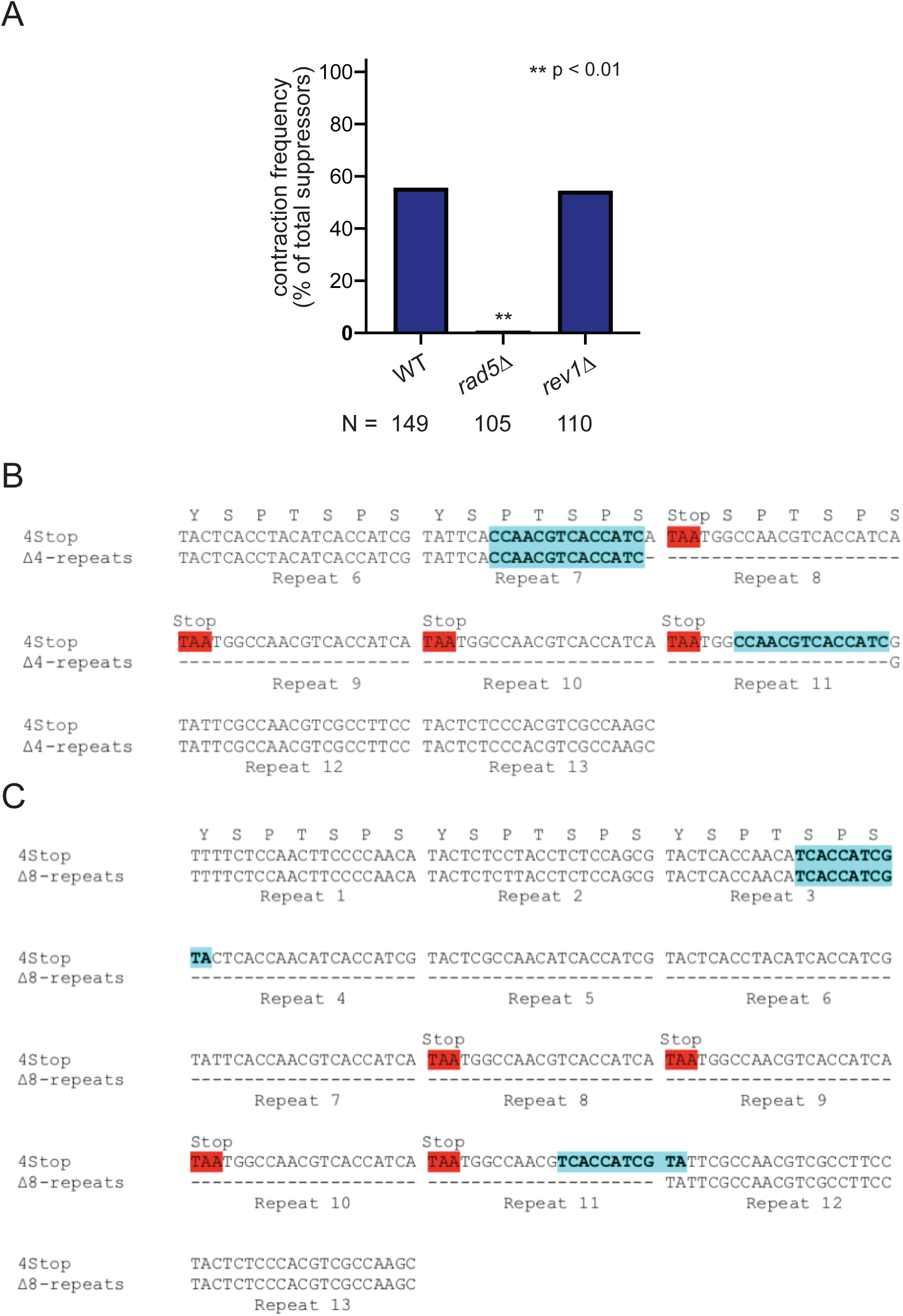
Template switching mediates contractions of the CTD. A) Contraction frequencies of the CTD in the absence of DNA repair proteins involved in post-replication repair. Frequencies are determined by the accumulation of data from three independent plasmid transformations, and statistical significance was determined using a two-proportions z-test. B) Sequence analysis of a contraction event in which the four repeats containing stop codons were deleted. A 14-bp microhomology is present before and after the deletion. C) Sequence analysis of a contraction event in which eight total repeats, including those containing stop codons, were deleted. An 11-bp microhomology is present before and after the deletion.

### Microhomologies flank deleted sequences in contraction events

In addition to the DNA lesions that are readily bypassed by TLS, there are many other types of replication barriers, such as DNA secondary structures, DNA-bound by proteins, and transcriptional machinery when replicating long and highly transcribed genes. When the replication fork collides with obstacles that cannot be bypassed by TLS, cells employ the other PRR pathway, Template Switching, to avoid fork collapse (22). In addition to its role in TLS, Rad5p is important for the initiation of TS through its role in the polyubiquitination of PCNA (23, 24). Template switching has been demonstrated to occur via two mechanisms, fork regression and strand invasion, and is typically thought to be error-free because in both cases it involves bypassing the replication obstacle using a homologous template (22, 25, 26). In order for template switching to generate contractions during a strand invasion repair event, the nascent strand associated with the obstructed arm of the replication fork would have to invade the sister chromatid at the incorrect locus, resulting in a failure to replicate one or more repeats. During template switching by fork reversal, the two nascent strands would have to misalign in order to generate a contraction. The presence of microhomologies within the DNA would facilitate misalignment between the nascent strands as well as strand invasion at the incorrect locus. We therefore sequenced contraction events in order to determine if microhomologies are present at the repair junctions.

Sequence analysis of the repair junctions revealed the presence of microhomologies flanking the deleted regions in all contraction events, which result in the loss of four, five, six, or eight repeats. The most commonly observed contraction event is the in-frame deletion of only the four repeats containing stop codons, repeats 8–11, flanked by a microhomology of 14 bp. Less frequent events involve deletion of up to four additional repeats preceding those containing stop codons, resulting in the loss of up to eight total repeats. Microhomologies flanking contractions ranged in size from 11 bp to 14 bp, but no correlation was observed between microhomology length and the frequency of an observed contraction. Sequences from deletions of four and eight repeats are shown in Figures 4B and 4C, respectively.

The microhomologies bordering the repair junctions that we have observed in the CTD are also consistent with repair by microhomology-mediated end-joining (MMEJ). In order to determine if this pathway plays a role in contractions of the CTD, we measured the contraction frequency in *rad1*Δ and *rad10*Δ backgrounds (Supplemental Figure 2). Rad1p/Rad10p acts a complex to cleave the flaps generated during MMEJ (27, 28). In the absence of Rad1p and Rad10p, contractions significantly increased from 56 percent in the wild type to 79 and 83 percent, respectively.

## DISCUSSION

Repetitive sequences have generally been dismissed as inconsequential; however, it is becoming increasingly clear that repetitive regions can play important roles in modulating protein function. Repeat copy number in most repetitive coding regions often varies not only at the organismal level, but from generation to generation (29). When found in open reading frames, variable repeats can lead to variable phenotypes, and repetitive sequences throughout the genome may serve as hotspots for evolution. Trinucleotide repeat instability is well known to play a role both in disease and beneficial phenotypic variation, and the mechanisms by which these repeats expand and contract are well studied (7, 8, 11). While more complex tandem repeats are also known to be variable and to contribute to variable phenotypes (10, 14, 30), less is known about the molecular mechanisms that contribute to variation.

The CTD of RNA polymerase II is a fascinating model for exploring repeat copy number variation as the sequence acts as a scaffolding domain that is essential for transcription, and its amino acid sequence is likewise highly conserved across eukaryotes. Despite this, the coding sequence is both highly polymorphic and under strong purifying selection to retain function. Using the heptapeptide repeat of the CTD of RNA polymerase II as a model, we set out to identify the mechanisms by which complex tandem repeats promote variability and drive evolution. In addition to variable repeat copy number across organisms, with humans having twice as many repeats as yeast, there is significant evidence of rearrangements of the CTD across strains of yeast (Figure 1). We previously determined that expansions of the CTD require Rad52p (4); in this work, we sought to determine the mechanism by which the CTD undergoes contractions.

In a *ku70*Δ background, the contraction frequency was not significantly different from wild type, suggesting that contractions do not occur by non-homologous end-joining. The contraction frequency was significantly increased in *pol32*Δ, *rad52*Δ, and *rad51*Δ backgrounds, indicating that contractions are not a result of break-induced replication or homologous recombination. In fact, our data support that homologous recombination is a competing mechanism for dealing with genetic instability. Ruling out the predominant DNA repair pathways, we hypothesized that PRR mediates contractions. Both PRR pathways (TS and TLS) require the ubiquitin ligase Rad5p. In a *rad5*Δ background, the frequency of contractions was reduced from 56% to less than 1%, indicating that contractions are mediated by one of the PRR pathways. TLS, which requires REV1, primarily results in point mutations and was not expected to lead to repeat instability. In a *rev1*Δ background, the contraction frequency of the CTD was not significantly different from wild type, confirming that TLS does not play a role in contractions of the CTD. Because contractions require Rad5p but are not mediated by TLS, we propose that contractions of the CTD occur by template switching. There are no known genetic factors that are involved exclusively in TS that can be isolated in order to directly probe the role of TS in contractions. However, we believe that the combination of the *rad5*Δ and *rev1*Δ mutants is sufficient to isolate TS.

Template switching allows cells to bypass replication obstacles such as base adducts, secondary structures, DNA-bound proteins, or transcriptional machinery in the case of highly transcribed genes. TS can occur by two mechanisms: 1) a strand invasion mechanism in which the nascent strand invades into the sister chromatid in order to bypass the obstacle and 2) a fork reversal mechanism in which the replication fork regresses, and the nascent strands dissociate from their respective templates and anneal to each other in order to bypass the obstacle (22). Template switching by strand invasion requires the RAD52 epistasis group (31, 32). Because we have demonstrated that Rad51p and Rad52p are not required for contractions of the CTD, our best understanding of the literature concludes that contractions are mediated by a fork regression event.

Repetitive sequences form several secondary structures known to impede DNA replication, such as hairpins, triplex DNA, and G-quadruplexes (19). While trinucleotide repeats readily form hairpins, the CTD consists of a degenerate 21-bp repeat whose structure-forming ability is not immediately apparent. Capra et al. previously reported G_4_-DNA near the 3’ end of the *RPB1* coding sequence, which encodes the CTD (33). G_4_-DNA is characterized by GG, GGG, or GGGG motifs that are connected by short loops and can form G-quadruplex structures, which consist of stacked G-tetrads (34). Consistent with the results of Capra et al., we found that GG motifs are enriched on the noncoding strand of the CTD compared to the body of the *RPB1* gene (Figure 5). Sequence analysis of the complete *RPB1* gene using the QGRS Mapper also confirms that predicted G-quadruplex forming sequences are enriched in the CTD (34). Despite the fact that the CTD represents only 10% of the total length of RPB1, this region has 33% of the predicted G-quadruplex forming sequences (Figure 5). We also recently demonstrated that several segments of the CTD are capable of forming G-quadruplex structures *in vitro* (4). *In vivo*, these structures could block the replication fork and necessitate fork reversal. Notably, the enrichment of GG motifs and predicted G-quadruplex forming sequences are isolated to the CTD. The CTD sequence therefore promotes variability, while the body of the essential *RPB1* gene is strictly conserved. In addition to being able to form secondary structures, *RPB1* is a long gene, and increased transcription time may result in collisions between the replicative and transcriptional machineries.

**Figure 5.**
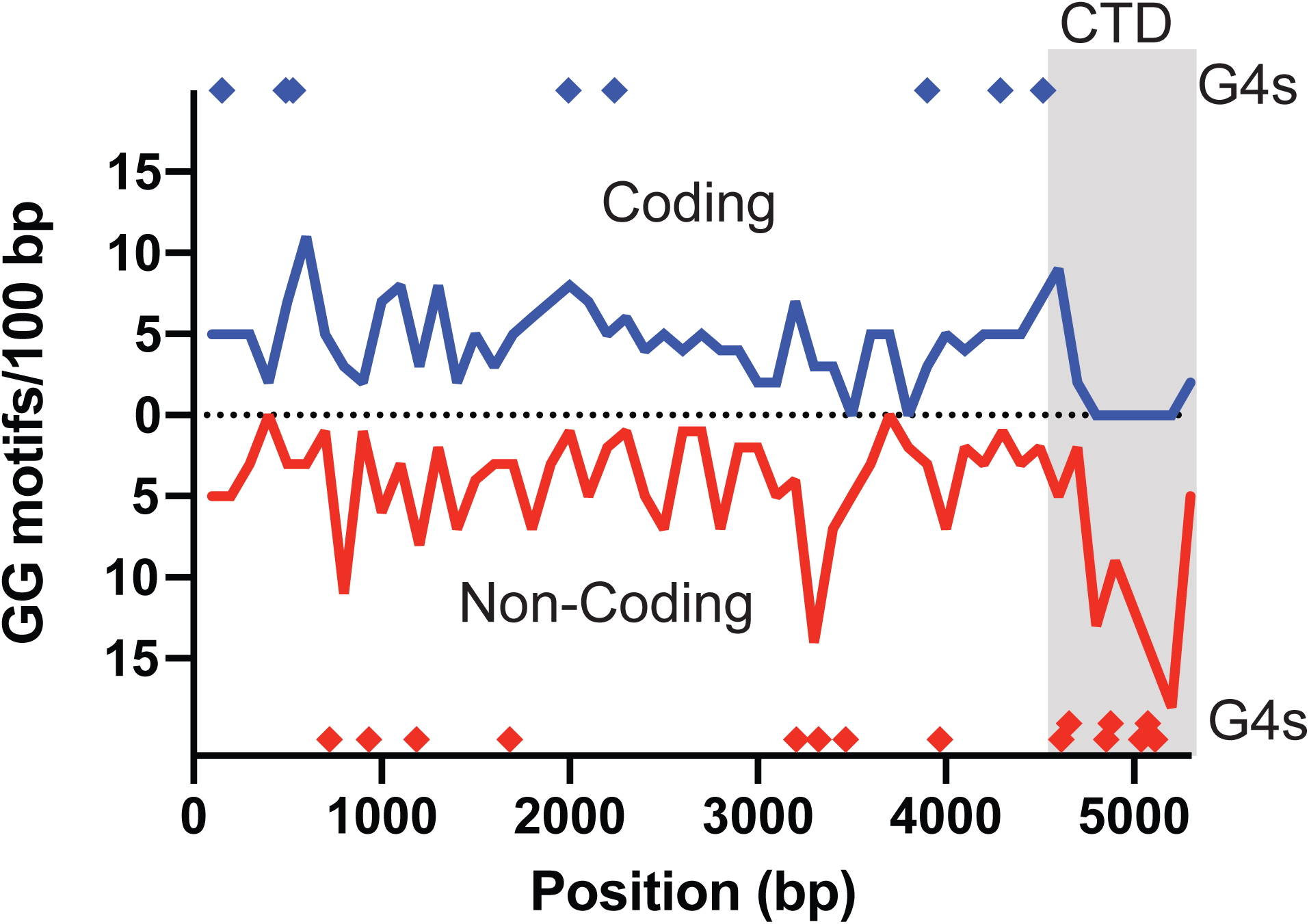
The CTD is enriched in predicted G-quadruplex-forming sequences. GG motifs on the coding (orange) and noncoding (blue) strands of *RPB1*. G-quadruplex-forming sequences predicted by the QGRS Mapper are highlighted. The region between 4624 and 5202 bp is the CTD.

When the replication fork encounters obstacles such as secondary structures and transcriptional machinery, it can be stabilized by fork reversal. Such template switching events are typically thought to be error-free due to the use of a homologous template as the source for bypassing the replication obstacle. Repetitive sequences are unusual, however, due to the presence of multiple homologous templates located in close proximity, which can result in repair events that change repeat copy number while maintaining the open reading frame and primary amino acid sequence. We examined the repair junctions of contraction events and discovered microhomologies ranging from 11 to 14 bp in length that could serve as sites of misalignment during template switching via fork reversal (Figure 4). This misalignment would result in a segment of the CTD looping out and being cleaved by an endonuclease, leading to a contraction (Figure 6A). The microhomologies present in the CTD could also serve as templates for repair by MMEJ. Unexpectedly, contractions increased in the absence of Rad1p and Rad10p, suggesting that contractions of the CTD do not require MMEJ. We are therefore confident in our model that contractions of the CTD are primarily mediated by fork reversal during template switching. This model is consistent with the results that contractions require Rad5p but not Rev1p, Rad52p, or Rad51p, indicating that contractions are mediated by a PRR pathway that does not require translesion synthesis or strand invasion. Combined with the presence of secondary structure-forming sequences in the CTD, this data supports a fork reversal mechanism, and the microhomologies that we have observed flanking contraction junctions provide clear evidence for how the two nascent strands can misalign, resulting in contractions of the CTD.

**Figure 6.**
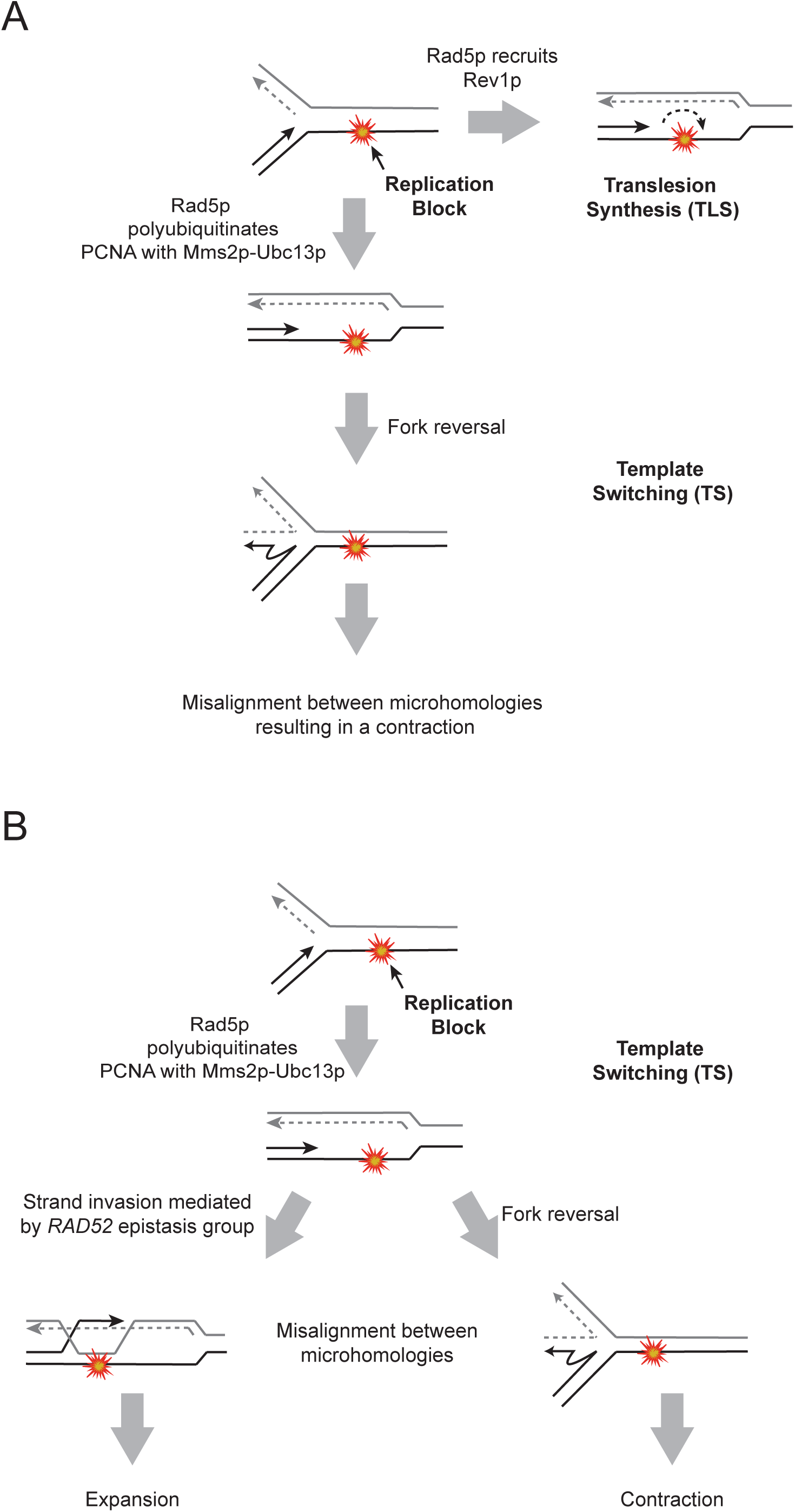
Template switching during post-replication repair mediates repeat instability of the CTD. A) During post-replication repair, the replication fork encounters an obstacle that it cannot bypass on its own. One repair option is TLS in which translesion polymerases replace the replicative polymerase to bypass the damage. Cells also employ template switching to bypass the obstacle using the sister chromatid as a template. Due to the presence of microhomologies throughout the CTD, we propose that misalignment during fork reversal leads to contractions of the CTD. B) When the replication fork encounters an obstacle, such as a G-quadruplex, template switching can be used to bypass it. TS is initiated by Rad5p-mediated polyubiquitination of PCNA and can occur by two mechanisms—strand invasion and fork reversal. During strand invasion, microhomologies in the CTD result in the re-replication of one or more repeats, leading to expansions. This process requires Rad52p. During fork reversal, microhomologies align, causing a segment of the CTD to loop out, resulting in a contraction.

Our previously published data indicates that expansions of the CTD require Rad52p (4). We therefore initially hypothesized that expansions are mediated by homologous recombination. Based on our new evidence that contractions of the CTD are mediated by template switching, we propose that like contractions, expansions of the CTD are also mediated by TS, but through a Rad52p-mediated strand invasion mechanism as opposed to a fork reversal mechanism (Figure 6B). This hypothesis is supported by the fact that expansions increased in a *pif1-m2* background (4). Pif1p is a DNA helicase that plays a role in unwinding G-quadruplex structures (35). In the absence of functional Pif1p, G-quadruplex structures formed by the CTD sequence may block the replication fork more frequently, promoting template switching.

Expansions and contractions of the CTD of RNA polymerase II are occurring all the time at a low frequency. This instability may serve as a mechanism to reduce mutagenesis in an essential sequence by allowing for the removal of damaged repeats while maintaining overall length. We have observed this phenomenon in our system through the removal of artificially introduced stop codons, and sequence analysis of 93 yeast strains from a wide range of sources demonstrates that this phenomenon occurs in wild populations as well (Figure 1 and Figure S1). The microhomologies that we observed flanking repair junctions provide clear evidence for how these rearrangements could have occurred. Our work demonstrates that the alignment of microhomologies during template switching events lead to variability in complex tandem repeats, enabling these sequences to both promote evolution and maintain functionality.

## SUPPLEMENTARY DATA

Supplementary Data is available from bioRxiv.

## ACKNOWLEDGEMENTS

The authors would like to acknowledge Summer Morrill, Mohammad Mosaheb, and Bradley Reinfeld for early contributions to this project. We also thank M. Parmenter, K. McElroy, and other members of the Fuchs Lab for their support and helpful discussions during the preparation of this manuscript.

