## Supplemental data for "Template Switching Mediates Contractions of a Repetitive Coding Region in *S. cerevisiae*"

#### SUPPLEMENTAL METHODS AND DATA

*Determining Mutation Rate of the Truncated CTD*— Fluctuation analysis was modified from Aksenova *et al.* to fit the DOX selection system (1). Briefly, strains expressing the 4stop construct were streaked for isolation on SC-Leu to maintain the plasmid and incubated at 30°C for 3-5 days, allowing colonies to grow to approximately ~$10^6$ cells (time necessary for sufficient growth varies by strain). For each experiment, 12 individual colonies were suspended in water and plated on SC-Leu+DOX; the same cell suspension was diluted $10^4$-fold and plated on YPD as a cell count control. After 3-5 days, colonies on both the DOX and control plates were counted, and colony counts were input into the fluctuation analysis calculator (FALCOR) in order to determine the mutation rate for each strain. FALCOR employs the MSS-maximum likelihood estimator method (MSS-MLE) (2). At least three independent fluctuation analysis experiments were performed for each strain, and the average rate of suppressor formation was reported.

**Supplemental Table 1. Primers used in this work**

| Description | Sequence 5' --> 3' |
|---|---|
| CTD reverse primer | TTTACTAGCGCCGTTGGTTT |
| CTD forward primer | GATCGATGAGGAGTCACTGG |
| Forward primer for amplification of the *ku70*Δ deletion | TGATTTGTTAAGTGACTCTAAGCCTGATTTTAAAACGGGAAT ATTATGCGTACGCTGCAGGTCGAC |
| Reverse primer for amplification of the *ku70*Δ deletion | ATATTGTATGTAACGTTATAGATATGAAGGATTTCAATCGTC TTTAATCGATGAATTCGAGCTCG |
| Forward primer for verification of the *ku70*Δ mutant | TTCAATGCAAAACAAAATTCCTT |
| Forward primer for amplification of the *rad52*Δ deletion | TTGCCAAGAACTGCTGAAGGTTCTGGTGGCTTTGGTGTGTT GTTGATGCGTACGCTGCAGGTCGAC |
| Reverse primer for amplification of the *rad52*Δ deletion | AAATAATGATGCAAATTTTTTATTTGTTTCGGCCAGGAAGCG TTTCAATCGATGAATTCGAGCTCG |
| Forward primer for verification of the *rad52*Δ mutant | AATGCAAACAAGGAGGTTGC |
| Forward primer for amplification of the *rad51*Δ deletion | CCAATCTAGTTTAGCTATCCTGCAA |
| Reverse primer for amplification of the *rad51*Δ deletion | AATTTTTCTCTTCACTCCCCTAAAA |
| Forward primer for amplification of the *pol32*Δ deletion | AACTAACAACCAGAAATAGGCTTTAGTTAACTCAATCGGTA ATTAATGCGTACGCTGCAGGTCGAC |
| Reverse primer for amplification of the *pol32*Δ deletion | ACATCACAATTAGTAATGGAAAGTGTTTGGAAAAAAAAGAA GATTAATCGATGAATTCGAGCTCG |
| Forward primer for verification of the *pol32*Δ mutant | ATGACGCCTTTTGATCCATT |
| Forward primer for amplification of the *rad5*Δ deletion | TCAAAAGGCCTTAGAAACACACCTAAAGTCTTACAGTATCA CAATATGCGTACGCTGCAGGTCGAC |
| Reverse primer for amplification of the *rad5*Δ deletion | AATAATAAATAAAGTCTTTATATATGAGTATGTGGTATGACT AATCGATGAATTCGAGCTCG |
| Forward primer for verification of the *rad5*Δ mutant | CTAAGCGCATTGCTCACTTG |
| Forward primer for amplification of the *rev1*Δ deletion | GCCGCGTTCACAGATTCCAA |
| Reverse primer for amplification of the *rev1*Δ deletion | CTATCGCTCATTTGGCTCTT |
| Reverse primer for verification of the *rev1*Δ mutant | CGAAGGGTGATTATCCGGTG |
| Forward primer for amplification of the *rad1*Δ deletion | GCCACAGTCAATATCGCGTC |
| Reverse primer for amplification of the *rad1*Δ deletion | TAATGCCACGCTCAGATTCC |
| Forward primer for verification of the *rad1*Δ mutant | TGATTCACTAATGCCACGCTC |

| | |
|---|---|
| Forward primer for amplification of the *rad10*Δ deletion | TCAGCTGCTCGGGATTTAGT |
| Reverse primer for amplification of the *rad10*Δ deletion | ATGGCGTGGAGGTGAGATAC |
| Forward primer for verification of the *rad10*Δ mutant | CGCAACCTGAATCGTGATGA |
| NATMX reverse primer for verification of the *rad52*Δ, *ku70*Δ, *pol32*Δ, and *rad5*Δ mutants | CGAGTACGAGATGACCACGA |
| KANMX forward primer for verification of the *rev1*Δ, *rad1*Δ, and *rad10*Δ mutants | TGATTTTGATGACGAGCGTAAT |

**Supplemental Table 2. Overall rates of suppressor formation in cells expressing p4stop determined by fluctuation analysis**

| Strain | Rate of Suppressor Formation ($x10^{-6}$ / cell / generation) |
|---|---|
| Wild type | 26.39 (10.9-37.9) |
| *ku70*Δ | 5.75 (2.4-8.6) |
| *rad52*Δ | 3.6 (2.5-4.7) |
| *rad51*Δ | 1.58 (2.5-6.6) |
| *pol32*Δ | 21.06 (7.3-43.0) |
| *rad5*Δ | 4.72 (3.9-5.2) |

**Supplemental Figure 1:** Alignment of CTD coding regions from *S. cerevisiae*. Strains with identical CTD coding sequences were grouped and give a letter with the number in parentheses referring to the number of sequences in a group. 24 unique sequences from 93 strains (3) were aligned using MAFFT and hand-corrected for errors in aligning repetitive sequences. Strains show multiple, genetically independent contraction and expansion events within the CTD coding region. Disagreements are highlighted with color.

**Supplemental Figure 2: Contraction frequencies of the CTD in the absence of DNA repair proteins involved in microhomology-mediated end-joining.** Frequencies are determined by the accumulation of data from three independent experiments, and statistical significance was determined using a two-proportions z-test.

### Figure S1

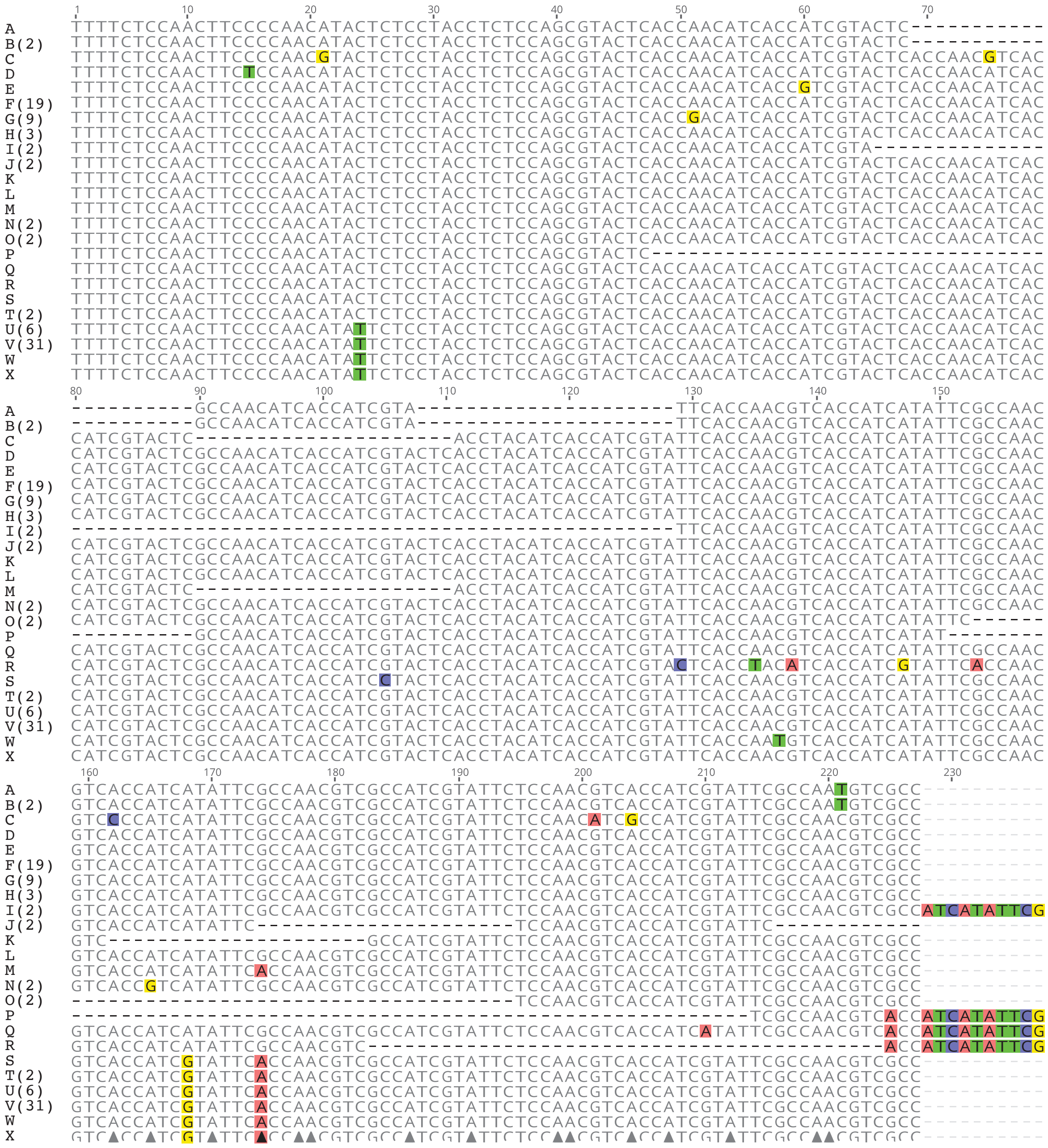

### Figure S1

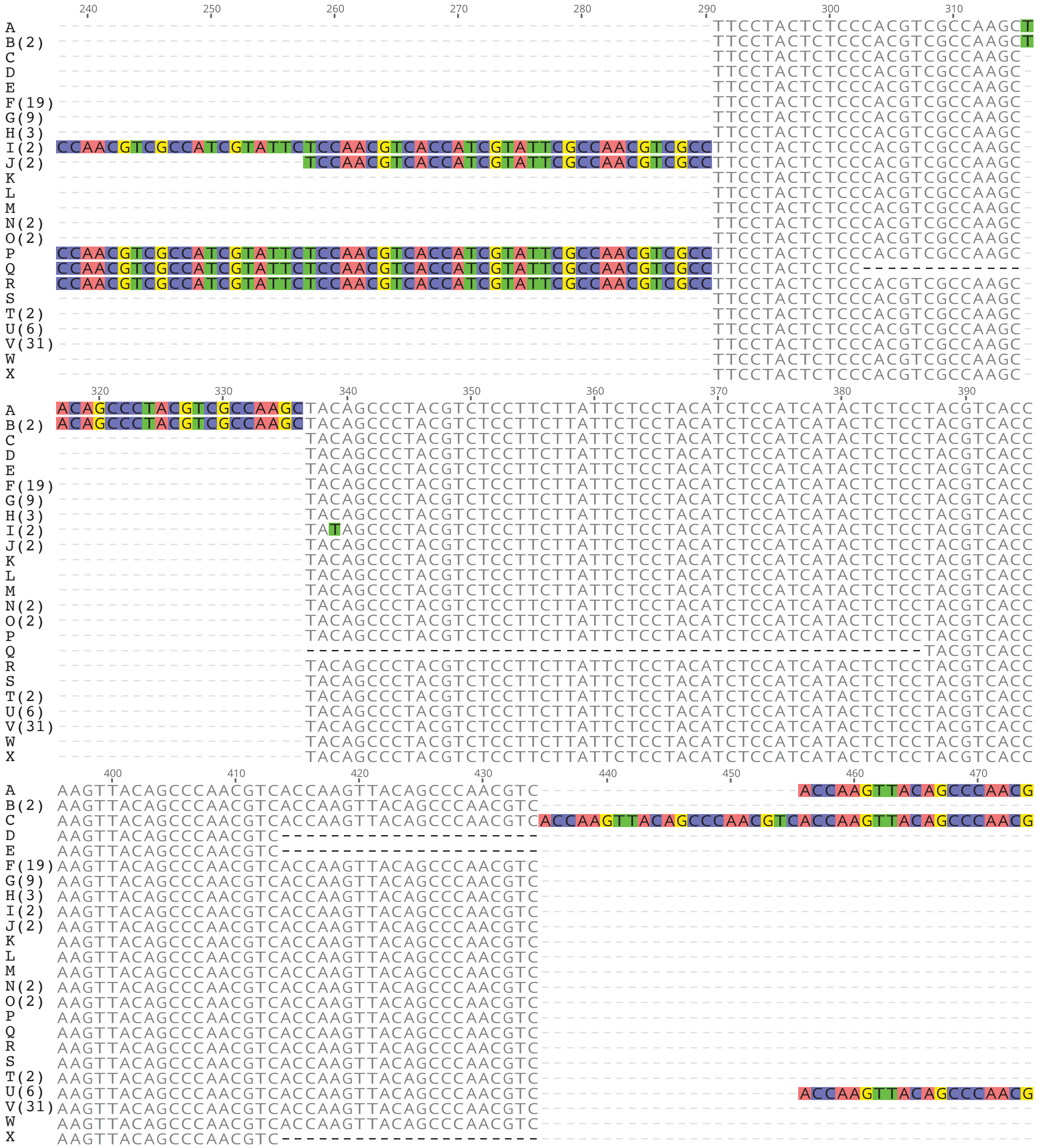

240
250
260
270
280
290
300
310
320
330
340
350
360
370
380
390
400
410
420
430
440
450
460
470
A
B(2)
C
D
E
F(19)
G(9)
H(3)
I(2)
J(2)
K
L
M
N(2)
O(2)
P
Q
R
S
T(2)
U(6)
V(31)
W
X

### Figure S1

```
A      TCTCCAGCCTATTCCCCAACATCACCAAGTTATAGTCCTACATCGCCTTCATACTCTCCAACGTCACCATCCTATTCCC
B(2)   --TCCAGCCTATTCCCCAACATCACCAAGTTATAGTCCTACATCGCCTTCATACTCTCCAACGTCACCATCCTATTCCC
C      TCTCCAGCCTATTCCCCAACATCACCAAGTTATAGTCCTACATCGCCTTCATACTCTCCAACGTCACCATCCTATTCCC
D      --TCCAGCCTATTCCCCAACATCACCAAGTTATAGTCCTACATCGCCTTCATACTCTCCAACGTCACCATCCTATTCCC
E      --TCCAGCCTATTCCCCAACATCACCAAGTTATAGTCCTACATCGCCTTCATACTCTCCAACATCACCATCCTATTCCC
F(19)  --TCCAGCCTATTCCCCAACATCACCAAGTTATAGTCCTACATCGCCTTCATACTCTCCAACATCACCATCCTATTCCC
G(9)   --TCCAGCCTATTCCCCAACATCACCAAGTTATAGTCCTACATCGCCTTCGTACTCTCCAACATCACCATCCTATTCCC
H(3)   --TCCAGCCTATTCCCCAACATCACCAAGTTATAGTCCTACATCGCCTTCATACTCTCCAACGTCACCATCCTATTCCC
I(2)   --TCCAGCCTATTCCCCAACATCACCAAGTTATAGTCCTACATCGCCTTCATACTCTCCAACATCACCATCCTATTCCC
J(2)   --TCCAGCCTATTCCCCAACATCACCAAGTTATAGTCCTACATCGCCTTCATACTCTCCAACATCACCATCCTATTCCC
K      --TCCAGCCTATTCCCCAACATCACCAAGTTATAGTCCTACATCGCCTTCATACTCTCCAACGTCACCATCCTATTCCC
L      --TCCAGCCTATTCCCCAACATCACCAAGTTATAGTCCTACATCGCCTTCATACTCTCCAACATCACCATCCTATTCCG
M      --TCCAGCCTATTCCCCAACATCACCAAGTTATAGTCCTACATCGCCTTCATACTCTCCAACATCACCATCCTATTCCC
N(2)   --TCCAGCCTATTCCCCAACATCACCAAGTTATAGTCCTACATCGCCTTCATACTCTCCAACATCACCATCCTATTCCC
O(2)   --TCCAGCCTATTCCCCAACATCACCAAGTTATAGTCCTACATCGCCTTCATACTCTCCAACATCACCATCCTATTCCC
P      --TCCAGCCTATTCCCCAACATCACCAAGTTATAGTCCTACATCGCCTTCATACTCTCCAACATCACCATCCTATTCCC
Q      --TCCAGCCTATTCCCCAACATCACCAAGTTATAGTCCTACATCGCCTTCATACTCTCCAACATCACCATCCTATTCCC
R      --TCCAGCCTATTCCCCAACATCACCAAGTTATAGTCCTACATCGCCTTCATACTCTCCAACATCACCATCCTATTCCC
S      --TCCAGCCTATTCCCCAACATCACCAAGTTATAGTCCTACATCGCCTTCATACTCTCCAACGTCACCATCCTATTCCC
T(2)   --TCCAGCCTATTCCCCAACATCACCAAGTTATAGTCCTACATCGCCTTCATACTCTCCAACGTCACCATCCTATTCCC
U(6)   TCTCCAGCCTATTCCCCAACATCACCAAGTTATAGTCCTACATCGCCTTCATACTCTCCAACGTCACCATCCTATTCCC
V(31)  --TCCAGCCTATTCCCCAACATCACCAAGTTATAGTCCTACATCGCCTTCATACTCTCCAACGTCACCATCCTATTCCC
W      --TCCAGCCTATTCCCCAACATCACCAAGTTATAGTCCTACATCGCCTTCATACTCTCCAACGTCACCATCCTATTCCC
X      --TCCAGCCTATTCCCCAACATCACCAAGTTATAGTCCTACATCGCCTTCATACTCTCCAACGTCACCATCCTATTCCC

                 560       570       580       590       600       610       620       630
A      CAACATCACCTTCTTACTCTCCCACCTCTCCAAACTATAGCCCTACTTCACCTTCTTACTCCCCAACATCTCCAGGCTA
B(2)   CAACATCACCTTCTTACTCTCCCACCTCTCCAAACTATAGCCCTACTTCACCTTCTTACTCCCCAACATCTCCAGGCTA
C      CAACATCACCTTCTTACTCTCCCACCTCTCCAAACTATAGCCCTACTTCACCTTCTTACTCCCCAACATCTCCAGGCTA
D      CAACATCACCTTCTTACTCTCCCACCTCTCCAAACTATAGCCCTACTTCACCTTCTTACTCCCCAACATCTCCAGGCTA
E      CAACATCACCTTCTTACTCTCCCACCTCTCCAAACTATAGCCCTACTTCACCTTCTTACTCCCCAACATCTCCAGGCTA
F(19)  CAACATCACCTTCTTACTCTCCCACCTCTCCAAACTATAGCCCTACTTCACCTTCTTACTCCCCAACATCTCCAGGCTA
G(9)   CAACATCACCTTCTTACTCTCCCACCTCTCCAAACTATAGCCCTACTTCACCTTCTTACTCCCCAACATCTCCAGGCTA
H(3)   CAACATCACCTTCTTACTCTCCCACCTCTCCAAACTATAGCCCTACTTCACCTTCTTACTCCCCAACATCTCCAGGCTA
I(2)   CAACATCACCTTCTTACTCTCCCACCTCTCCAAACTATAGCCCTACTTCACCTTCTTACTCCCCAACATCTCCAGGCTA
J(2)   CAACATCACCTTCTTACTCTCCCACCTCTCCAAACTATAGCCCTACTTCACCTTCTTACTCCCCAACATCTCCAGGCTA
K      CAACATCACCTTCTTACTCTCCCACCTCTCCAAACTATAGCCCTACTTCACCTTCTTACTCCCCAACATCTCCAGGCTA
L      CAACATCACCTTCTTACTCTCCCACCTCTCCAAACTATAGCCCTACTTCACCTTCTTACTCCCCAACATCTCCAGGCTA
M      CAACATCACCTTCTTACTCTCCCACCTCTCCAAACTATAGCCCTACTTCACCTTCTTACTCCCCAACATCTCCAGGCTA
N(2)   CAACATCACCTTCTTACTCTCCCACCTCTCCAAACTATAGCCCTACTTCACCTTCTTACTCCCCAACATCTCCAGGCTA
O(2)   CAACATCACCTTCTTACTCTCCCACCTCTCCAAACTATAGCCCTACTTCACCTTCTTACTCCCCAACATCTCCAGGCTA
P      CAACATCACCTTCTTACTCTCCCACCTCTCCAAACTATAGCCCTACTTCACCTTCTTACTCCCCAACATCTCCAGGCTA
Q      CAACATCACCTTCTTACTCTCCCACCTCTCCAAACTATAGCCCTACTTCACCTTCTTACTCCCCAACATCTCCAGGCTA
R      CAACATCACCTTCTTACTCTCCCACCTCTCCAAACTATAGCCCTACTTCACCTTCTTACTCCCCAACATCTCCAGGCTA
S      CAACATCACCTTCTTACTCTCCCACCTCTCCAAACTATAGCCCTACTTCACCTTCTTACTCCCCAACATCTCCAGGCTA
T(2)   CAACATCACCTTCTTACTCTCCCACCTCTCCAAACTATAGCCCTACTTCACCTTCTTACTCCCCAACATCTCCAGGCTA
U(6)   CAACATCACCTTCTTACTCTCCCACCTCTCCAAACTATAGCCCTACTTCACCTTCTTACTCCCCAACATCTCCAGGTTA
V(31)  CAACATCACCTTCTTACTCTCCCACCTCTCCAAACTATAGCCCTACTTCACCTTCTTACTCCCCAACATCTCCAGGCTA
W      CAACATCACCTTCTTACTCTCCCACCTCTCCAAACTATAGCCCTACTTCACCTTCTTACTCCCCAACATCTCCAGGCTA
X      CAACATCACCTTCTTACTCTCCCACCTCTCCAAACTATAGCCCTACTTCACCTTCTTACTCCCCAACATCTCCAGGCTA

                  640       650       660       670       680       690       700  705
A      CAGCCCAGGATCTCCTGCATATTCTCCAAAGCAAGACGAACAAAAGCATAATGAAAATGAAAATTCCAGATGA
B(2)   CAGCCCAGGATCTCCTGCATATTCTCCAAAGCAAGACGAACAAAAGCATAATGAAAATGAAAATTCCAGATGA
C      CAGCCCAGGATCTCCTGCATATTCTCCAAAGCAAGACGAACAAAAGCATAATGAAAATGAAAATTCCAGATGA
D      CAGCCCAGGATCTCCTGCATATTCTCCAAAGCAAGACGAACAAAAGCATAATGAAAATGAAAATTCCAGATGA
E      CAGCCCAGGATCTCCTGCATATTCTCCAAAGCAAGACGAACAAAAGCATAATGAAAATGAAAATTCCAGATGA
F(19)  CAGCCCAGGATCTCCTGCATATTCTCCAAAGCAAGACGAACAAAAGCATAATGAAAATGAAAATTCCAGATGA
G(9)   CAGCCCAGGATCTCCTGCATATTCTCCAAAGCAAGACGAACAAAAGCATAATGAAAATGAAAATTCCAGATGA
H(3)   CAGCCCAGGATCTCCTGCATATTCTCCAAAGCAAGACGAACAAAAGCATAATGAAAATGAAAATTCCAGATGA
I(2)   CAGCCCAGGATCTCCTGCATATTCTCCAAAGCAAGACGAACAAAAGCATAATGAAAATGAAAATTCCAGATGA
J(2)   CAGCCCAGGATCTCCTGCATATTCTCCAAAGCAAGACGAACAAAAGCATAATGAAAATGAAAATTCCAGATGA
K      CAGCCCAGGATCTCCTGCATATTCTCCAAAGCAAGACGAACAAAAGCATAATGAAAATGAAAATTCCAGATGA
L      CAGCCCAGGATCTCCTGCATATTCTCCAAAGCAAGACGAACAAAAGCATAATGAAAATGAAAATTCCAGATGA
M      CAGCCCAGGATCTCCTGCATATTCTCCAAAGCAAGACGAACAAAAGCATAATGAAAATGAAAATTCCAGATGA
N(2)   CAGCCCAGGATCTCCTGCATATTCTCCAAAGCAAGACGAACAAAAGCATAATGAAAATGAAAATTCCAGATGA
O(2)   CAGCCCAGGATCTCCTGCATATTCTCCAAAGCAAGACGAACAAAAGCATAATGAAAATGAAAATTCCAGATGA
P      CAGCCCAGGATCTCCTGCATATTCTCCAAAGCAAGACGAACAAAAGCATAATGAAAATGAAAATTCCAGATGA
Q      CAGCCCAGGATCTCCTGCATATTCTCCAAAGCAAGACGAACAAAAGCATAATGAAAATGAAAATTCCAGATGA
R      CAGCCCAGGATCTCCTGCATATTCTCCAAAGCAAGACGAACAAAAGCATAATGAAAATGAAAATTCCAGATGA
S      CAGCCCAGGATCTCCTGCATATTCTCCAAAGCAAGACGAACAAAAGCATAATGAAAATGAAAATTCCAGATGA
T(2)   CAGCCCAGGATCTCCTGCATATTCTCCAAAGCAAGACGAACAAAAGCATAATGAAAATGAAAATTCCAGATGA
U(6)   CAGCCCAGGATCTCCTGCATATTCTCCAAAGCAAGACGAACAAAAGCATAATGAAAATGAAAATTCCAGATGA
V(31)  CAGCCCAGGATCTCCTGCATATTCTCCAAAGCAAGACGAACAAAAGCATAATGAAAATGAAAATTCCAGATGA
W      CAGCCCAGGATCTCCTGCATATTCTCCAAAGCAAGACGAACAAAAGCATAATGAAAATGAAAATTCCAGATGA
X      CAGCCCAGGATCTCCTGCATATTCTCCAAAGCAAGACGAACAAAAGCATAATGAAAATGAAAATTCCAGATGA
```

Figure S2

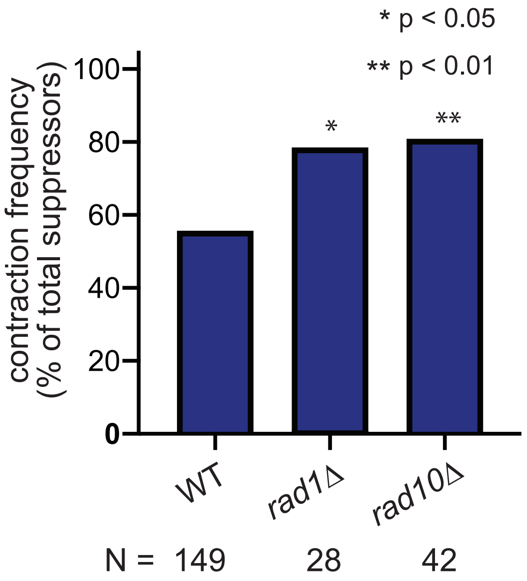

* p < 0.05
** p < 0.01
contraction frequency
(% of total suppressors)
100
80
60
40
20
0
*
**
WT
rad1Δ
rad10Δ
N = 149 28 42
